# Identification of essential genes for *Escherichia coli* aryl polyene biosynthesis and function in biofilm formation

**DOI:** 10.1101/2020.04.22.055939

**Authors:** Isabel Johnston, Lucas J Osborn, Rachel L Markley, Elizabeth A McManus, Anagha Kadam, Karlee B Schultz, Nagashreyaa Nagajothi, Philip P Ahern, J Mark Brown, Jan Claesen

**Affiliations:** Department of Cardiovascular and Metabolic Sciences, Lerner Research Institute, Cleveland Clinic, Cleveland, OH, USA; Department of Molecular Medicine, Cleveland Clinic Lerner College of Medicine of Case Western Reserve University, Cleveland, OH, USA; National Cancer Institute, National Institutes of Health, Bethesda, MD, USA; College of Arts and Sciences, John Carroll University, University Heights, OH, USA; University Honors College, University of Pittsburgh, Pittsburgh, PA, USA; Center for Microbiome and Human Health, Cleveland Clinic, Cleveland, OH, USA

**Author notes:** LJO and RLM contributed equally to this work.

## Abstract

Aryl polyenes (APEs) are specialized polyunsaturated carboxylic acids that were identified *in silico* as the product of the most widespread family of bacterial biosynthetic gene clusters (BGCs). They are present in several Gram-negative host-associated bacteria, including multi-drug resistant human pathogens. Here, we characterize a biological function of APEs, focusing on the BGC from a uropathogenic *Escherichia coli* (UPEC) strain. We first perform a genetic deletion analysis to identify the essential genes required for APE biosynthesis. Next, we show that APEs function as fitness factors that increase protection from oxidative stress and contribute to biofilm formation. Together, our study highlights key steps in the APE biosynthesis pathway that can be explored as potential drug targets for complementary strategies to reduce fitness and prevent biofilm formation of multi-drug resistant pathogens.

## INTRODUCTION

In a global search for novel types of bacterial secondary metabolites, we previously identified aryl polyenes (APEs) as the products of the most abundant biosynthetic gene cluster (BGC) family [1]. APEs are present in most major bacterial genera throughout the Proteobacteria and Bacteroidetes. They are often found in host-associated bacteria, including commensals as well as pathogens of humans, animals, and plants [1]. Among these are several human pathogens that are a major concern in nosocomial multidrug-resistant infections, including *Acinetobacter baumannii, Burkholderia cenocepacia*, as well as several *Enterobacter* isolates [2, 3]. APE BGCs are also present in most types of pathogenic *E. coli*, whereas they are typically absent from commensal and laboratory strains of *E. coli* [1].

APEs produced by diverse bacterial genera share a remarkably similar chemical scaffold, consisting of an aryl head group conjugated to a polyene carboxylic acid tail [1, 4–7] (Fig. 1). The main differences between APEs of different bacterial genera are in the polyene chain length and in the hydroxylation, methylation, or halogenation of the head group. This structural similarity is reflected in the observation that phylogenetically diverse APE BGCs share conserved core gene functions, resembling biosynthesis machinery for fatty acids and type II polyketides. Notably, an *in silico* comparison between BGCs revealed differential enzymatic functions in genus-dependent mechanisms for biosynthesis initiation, cell envelope attachment, and head group tailoring. APE head groups can be tailored by methylation (as in the *E. coli* CFT073 APE_Ec_ and flexirubin [1, 8, 9]) and halogenation (as in xanthomonadin [10]), while the hydroxyl group is commonly derived from the corresponding precursor molecule. While the core APE metabolites are similar in structure, further investigations are required to determine whether APEs also share a conserved lipid anchor in the outer membrane. Heterologous expression of the BGCs from *E. coli* CFT073 (APE_Ec_) and *Vibrio fischeri* ES114 (APE_Vf_) confirmed that these BGCs encompass all genes required for APE biosynthesis [1]. Moreover, the biosynthesis of the core APE moiety from the nematode symbiont *Xenorhabdus doucetiae* DSM17909 (APE_Xd_) was recently reconstituted *in vitro* by Grammbitter *et al*. [11]. The functions of nine enzymes from the APE_Xd_ BGC were characterized after heterologous expression and purification from *E. coli* and used to form APE_Xd_ *in vitro*. Our current study addresses outstanding questions in APE biology, including the identity of the essential genes for *in vivo* biosynthesis and uncovering their biological function among the bacteria that produce them.

**Figure 1.**
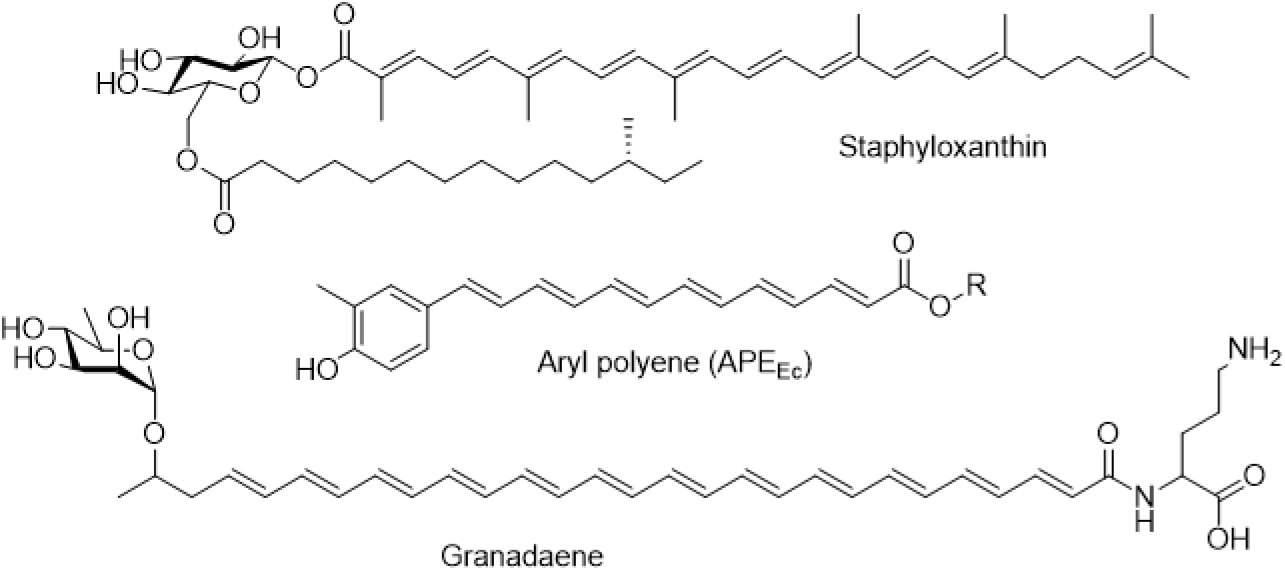
Chemical structures of pigmented fitness factors localized in the cell envelope of bacterial pathogens. Staphyloxanthin is the carotenoid produced by *S. aureus* responsible for the organism’s characteristic golden color (top). Aryl polyenes are widespread yellow pigments among Gram-negative pathogens (middle), for example APE_Ec_ from uropathogenic *E. coli*. Granadaene is an ornithine rhamno-polyene that constitutes the red pigmentation in Group B *Streptococcus* (bottom) [1, 16, 17].

APEs are covalently attached in the Gram-negative outer membrane [12], allowing the bacteria that produce them to potentially alter interactions with the surrounding environment. Among the best studied bacterial lifestyle changes that involve alterations in the expression of cell envelope structures are aggregation, adhesion to surfaces, and biofilm formation. Biofilms form when bacteria switch from planktonic growth to an adherent, three-dimensional lifestyle by embedding themselves in a self-produced extracellular matrix [13]. Biofilm formation impacts bacterial physiology, affecting pathogenesis and antibiotic resistance. Decreased antibiotic susceptibility can be due to a slowing of bacterial growth and metabolic activity, as well as reduced ability of antibiotic molecules to penetrate the biofilm matrix and reach interior cells [14, 15]. Moreover, growth in biofilms is associated with horizontal gene transfer, leading to more rapid strain adaptation and evolution. Since pathogens are more resilient when growing in biofilms, understanding the underlying mechanisms of biofilm formation and devising strategies to prevent and eliminate this process represent an important approach to limiting their pathogenic effects.

Gram-positive bacterial pathogens attach different types of polyenes in their cell envelope, including staphyloxanthin and granadaene (Fig. 1), which are polyenic virulence factors localized in the cell envelope of *Staphylococcus aureus* and Group B *Streptococcus*, respectively [16, 17]. Staphyloxanthin is a ‘golden’ carotenoid pigment produced from the mevalonate pathway, and granadaene is a red ornithine rhamno-polyene produced by fatty acid-like biosynthetic machinery [16, 17]. Despite their different biosynthetic origins, both of these compounds share remarkable analogous functions: they protect their producers from damage induced by reactive oxygen species (ROS) [18, 19]. Exogenous ROS can originate from a variety of sources, such as UV light, certain antibiotics or the host immune system. Neutrophils and macrophages utilize ROS and reactive nitrogen species (RNS) production as an initial line of defense against invading bacterial pathogens [20]. The lethal effect of ROS can be attributed to widespread damage to bacterial cells via induction of DNA double strand breaks and oxidation of proteins and lipids [21, 22]. Polyenes that are localized in the cell envelope can scavenge external ROS, thereby preventing more extensive damage to essential cellular molecules and increasing bacterial fitness. Hence, characterization of the staphyloxanthin biosynthetic pathway has fueled the development of an anti-virulence strategy for managing *S. aureus* infections [23].

In this study, we focus on the functional characterization of the APE_Ec_ BGC from uropathogenic *E. coli*. Using a heterologous expression and gene deletion approach, we identify the genes in the BGC that are required for APE_Ec_ biosynthesis. Bacterial strains that express APE_Ec_ are conferred with a fitness advantage compared to their APE^-^ counterparts. APE_Ec_ producers have an increased viability when subjected to an acute oxidative stress challenge and have an increased capability to form biofilms. Our work marks an important step in the biological understanding of APE_Ec_ function and paves the way for the development of complementary strategies to reduce *in vivo* fitness and biofilm formation of multidrug-resistant human pathogens.

## RESULTS

### The *E. coli* APE_Ec_ BGC contains predicted redundant functions and a dedicated transport system

We previously identified APEs as the product of one of the most abundant bacterial BGC families and chose the BGC from *E. coli* CFT073 for in-depth characterization. Heterologous expression of this 18 gene, 15.5 kb BGC showed that this region contains all the genes required for production of APE_Ec_ (Fig. 2A, [1]). All genes in the cluster are in the same orientation and several have overlapping open reading frames, suggesting the cluster is transcribed as one or multiple polycistronic mRNAs. The APE_Ec_ BGC encodes some seemingly duplicated functions, for example there are two acyl carrier proteins (ACPs) (ApeE and F), two 3-hydroxyacyl-ACP dehydratases (ApeI and P) and three 3-ketoacyl-ACP synthases (ApeC, O and R, Fig. 2, Table 1). *In vitro* studies by Grammbitter *et al*. [11] on the APE_Xd_ BGC showed that ApeR is required for initiation and that ApeC functions as a chain length factor through heterocomplex formation with ApeO, which is responsible for chain elongation. Interestingly, the ApeC chain length factor is encoded near the beginning of the cluster, whereas most of the other core biosynthesis genes are located quite distantly, near the cluster’s end.

**Figure 2.**
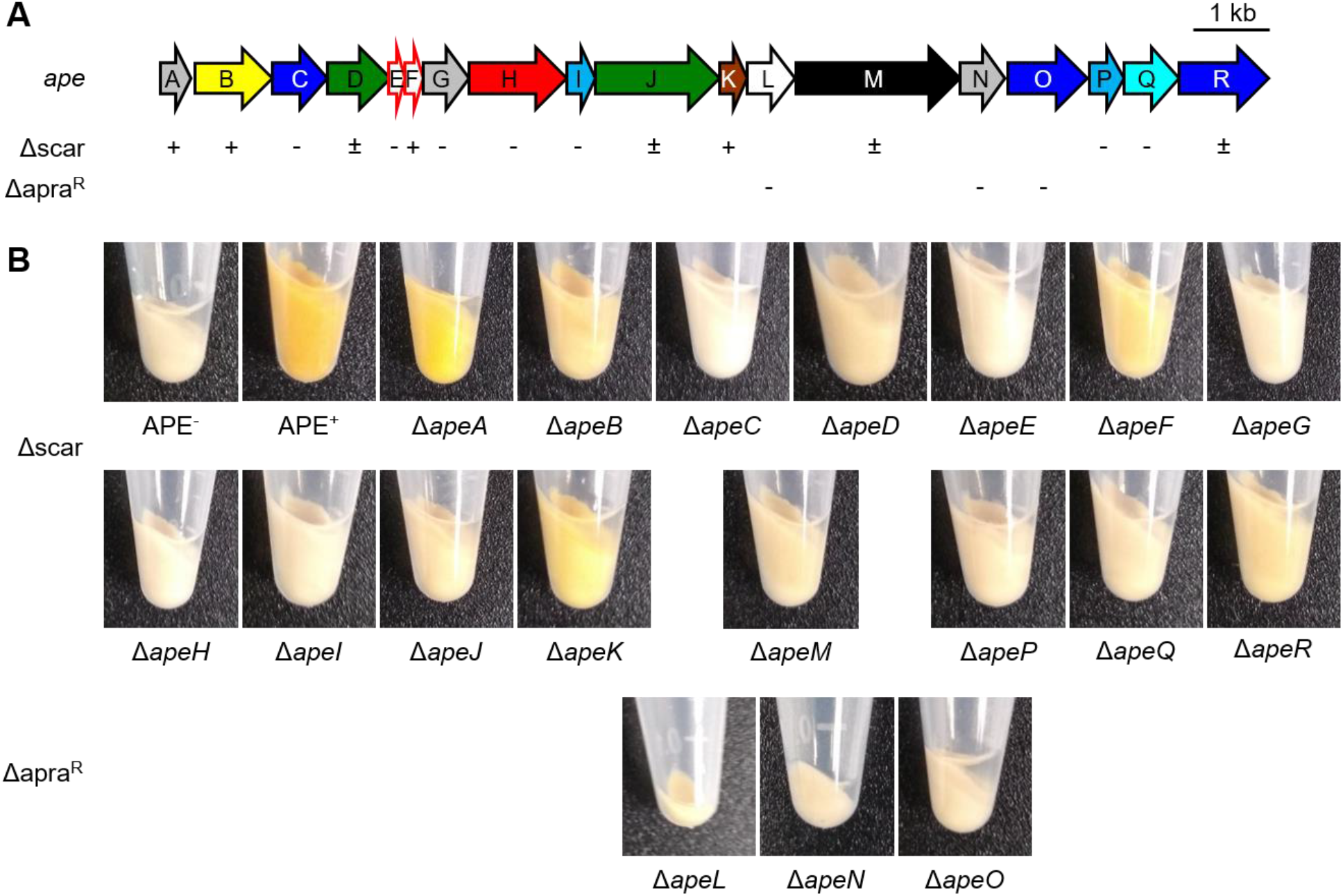
Pigmentation phenotype of individual deletion mutants in the APE_Ec_ BGC. **A)** Schematic arrow representation of the *E. coli* CFT073 APE_Ec_ BGC (see Table 1 for predicted gene functions). The pigmentation phenotype of individual scarred deletion mutants (Δscar) is reported below the BGC as follows: + (near wild-type level pigmentation), ± (largely reduced pigmentation), – (non-pigmented). We did not obtain scarred deletion mutants for *apaL, apeN* and *ape*O, and their respective Δapra^R^ mutants are not pigmented. **B)** Pigmentation phenotypes of the cell pellets from individual Δ*ape* strains in comparison to cell pellet pigmentation from APE^-^ and APE^+^ cultures. The cell pellet extracts (reconstituted in MeOH) of these strains are depicted in Fig. S2.

**Table 1.** Predicted gene functions for the APE_Ec_ BGC.

| ***ape*** | **Locus tag** | **Predicted function** |
| --- | --- | --- |
| **A** | c1204 | lipoprotein |
| **B** | c1203 | SAM-dependent methyltransferase |
| **C** | c1202 | 3-ketoacyl-ACP synthase |
| **D** | c1201 | lysophospholipid acyltransferase |
| **E** | c1200 | acyl carrier protein |
| **F** | c1199 | acyl carrier protein |
| **G** | c1198 | COG4648 - pyrophosphatase |
| **H** | c1197 | acyl-ACP synthetase |
| **I** | c1196 | 3-hydroxyacyl-ACP dehydratase |
| **J** | c1194 | glycosyltransferase / acyltransferase |
| **K** | c1193 | thioesterase |
| **L** | c1192 | LolA-like molecular chaperone |
| **M** | c1191 | RND family transporter |
| **N** | c1190 | DUF3261 – LolB-like membrane protein |
| **O** | c1189 | 3-ketoacyl-ACP synthase |
| **P** | c1188 | 3-hydroxyacyl-ACP dehydratase |
| **Q** | c1187 | 3-ketoacyl-ACP reductase |
| **R** | c1186 | 3-ketoacyl-ACP synthase |

The APE_Ec_ biosynthetic pathway is predicted to start with the activation of 4-hydroxybenzoic acid (4-HBA) by the specialized acyl-ACP synthetase ApeH (Fig. 3). *In vitro* reconstitution of the ApeH activity from the APE_Xd_ BGC revealed that this enzyme is capable of 4-HBA adenylation and ACP-loading without the formation of an intermediate CoA-thioester [11]. In *E. coli* and other Gram-negative bacteria, 4-HBA is produced by chorismate lyase (UbiC) as the first step in the ubiquinone biosynthesis pathway [24]. The APE_Ec_ cluster does not contain a dedicated phosphopantetheinyl transferase (PPTase) to activate the ACPs, ApeEF. An orphan PPTase, AcpT, was previously identified in *E. coli* O157:H7 EDL933, and *in vitro* biochemical analysis showed it can activate the respective ApeEF homologues in that strain [25]. The aromatic starter is further elongated with malonyl-ACP units by repetitive action of core biosynthetic enzymes related to those involved in fatty-acid and type II polyketide biosynthesis (Fig. 3). Methylation of the APE_Ec_ aromatic head group is likely catalyzed by the S-adenosyl-L-methionine dependent methyltransferase ApeB. The exact timing of this methylation step has not yet been determined and as a result of *in vitro* analysis of the APE_Xd_ methyltransferase, Grammbitter *et al*. postulated that methylation occurs post-APE biosynthesis [11]. The putative thioesterase ApeK could be involved in assisting the release of the finished APE_Ec_-carboxylic acid from the ACP. Next, APE_Ec_ is predicted to be acylated to a larger molecular entity in the membrane by two acyltransferases (predicted glycerol-3-phosphate acyltransferase ApeD and glycosyltransferase ApeJ), possibly via a glycerophospholipid-derived intermediate. Finally, the APE-containing molecule is localized to the outer membrane via a dedicated transport system comprised of ApeM, a “resistance-nodulation-division” (RND) family transmembrane protein and ‘localization of lipoprotein’ (Lol) homologues ApeL and ApeN (Table 1, [12, 26, 27]).

**Figure 3.**
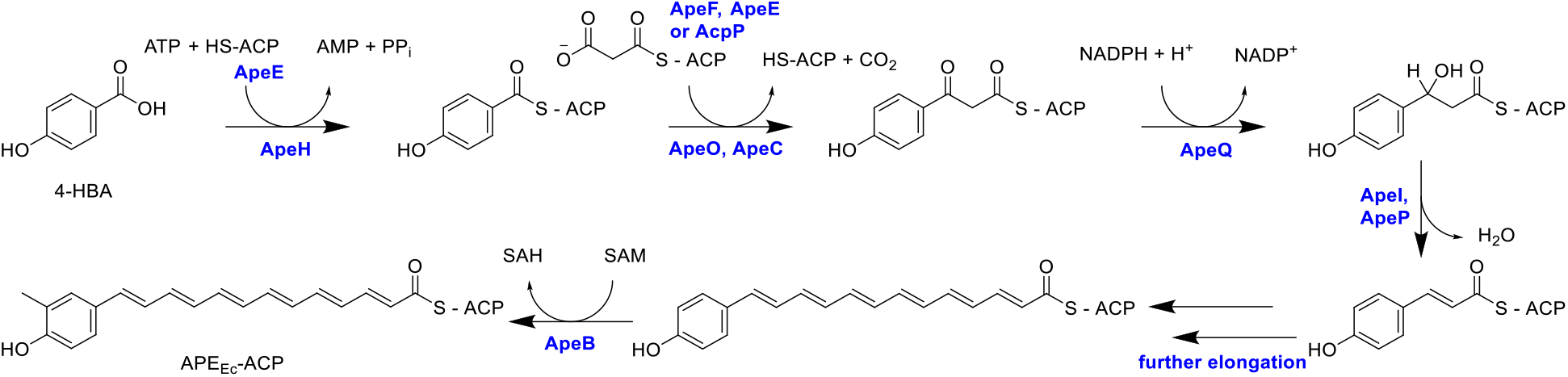
Proposed pathway for APE_Ec_ carboxylic acid biosynthesis. APE_Ec_ biosynthesis is predicted to start with the ACP-activation of 4-hydroxybenzoic acid (4-HBA), an early intermediate in ubiquinone biosynthesis. The aromatic starter is further elongated with malonyl-ACP units (potentially carried by ApeF, ApeE, or the ACP involved in fatty acid biosynthesis, AcpP [30]) by repetitive action of fatty-acid-like type II polyketide biosynthetic enzymes: ketosynthase ApeO and ketosynthase-chain length factor ApeC, ketoreductase ApeQ and dehydratases ApeI and/or ApeP. The SAM-dependent methyltransferase ApeB is involved in methylation of the aromatic head group, and it remains to be determined whether this modification occurs before or after ApeK (thioesterase) mediates release of the APE_Ec_-carboxylic acid from its ACP. Involvement of core enzymes in the pathway are indicated for *E. coli* CFT073 (in blue). SAM = *S*-adenosyl-L-methionine, SAH = *S*-adenosyl-L-homocysteine.

### Genetic analysis of the APE_Ec_ BGC reveals essential biosynthesis genes

We have constructed deletion mutants to link the individual genes in the APE_Ec_ BGC to their role in the biosynthetic pathway and identify potential functionally redundant genes *in vivo* (Table 1). To avoid potential polar effects of gene deletions within polycistronic operons, we used an in-frame “scarred” strategy involving a two-step PCR-generated recombineering procedure (Fig. S1, [28]). In the first step, the coding sequence between the start and stop codon for a gene of interest is replaced using a PCR cassette containing an antibiotic resistance marker with flanking FRT sites and homologous regions to the target gene. For overlapping genes, we designed in-frame deletion constructs that keep the entire overlapping open reading frame intact. In the second step, Flp recombinase is used to excise the resistance marker by recombining the FRT sites. In the resulting deletion mutant, the gene’s coding region is replaced by an in-frame 81 nucleotide “scar” sequence that lacks stop codons in all reading frames, thereby avoiding potential polar effects on downstream genes.

Upon heterologous introduction of the 15.5 kb APE_Ec_ BGC in a standard *E. coli* Top10 cloning strain (devoid of native BGC), we observed a strong yellow pigmentation phenotype [1]. Since APEs are covalently attached to the outer membrane [12], pigmentation is cell-associated and does not diffuse out into the culture medium. We used cell pellet pigmentation as a read-out for APE_Ec_ pathway functionality: mutant plasmids that conferred wild-type level pigmentation, comparable to the pJC121 containing parent strain are designated APE^+^; mutant plasmids that conferred reduced pigmentation are APE^±^; and mutant plasmids that did not display any pigmentation and thus were comparable to *E. coli* Top10 harboring the SuperCosI empty vector control were designated APE^-^ (Fig. 2A, B). As expected, most mutants in predicted core biosynthetic genes (represented in shades of blue in Fig. 2A) are APE^-^. Only deletion of *apeA*, the first gene in the BGC, retains wild type levels of pigmentation, suggesting it is not required for APE_Ec_ biosynthesis. Surprisingly, deletion of one of the two ACPs, Δ*apeF*, retains partial pigmentation. Both acyltransferase mutants, Δ*apeD* and Δ*apeJ*, retained partial pigmentation as well. The latter observation suggests the formation of intermediates in the cell envelope attachment process. We did not obtain scarred deletion mutants for *apeL, apeN* or *apeO* and the APE^-^ phenotype of their respective Δapra^R^ mutants could be due to polar effects on essential downstream genes in the BGC. In summary, individual deletions in most of the core biosynthesis and transport genes abolished APE-associated pigmentation (Table 1 and Fig. 2), whereas deletion of genes that are predicted to occur later in the pathway result in reduced pigmentation that could be caused by reduced flux through the pathway and improperly attached APE_Ec_ intermediates.

### Mutants that retain pigmentation produce biosynthetic or attachment intermediates

To characterize potential APE_Ec_ intermediates produced by our mutant strains, we performed a crude extraction of their cell pellets (Fig. S2) and profiled the APE-related metabolites in these extracts by high-performance liquid chromatography (HPLC; Figs. 4, S3). The primary APE_Ec_-lipid eluting near 100% ACN only seems to occur in the APE^+^ and Δ*apeR* extracts. The Δ*apeA*, Δ*apeB* and Δ*apeK* strains produce a comparable set of immature APE carboxylic acids that could be early intermediates formed before commencing the attachment pathway (Fig. 4). This was a remarkable observation provided the near wild-type levels of pigmentation in the Δ*apeA* strain. Our HPLC data indicate that *apeA* is not obsolete for full assembly of the APE_Ec_-lipid. Δ*apeD*, Δ*apeJ* and Δ*apeR* produce late pathway intermediates, which likely consist of APE-carboxylic acids that are impaired in their proper attachment to the final membrane lipid anchor. Deletion of the *apeF* ACP (orange) results in the formation of a series of yellow pigmented intermediates, thus indicating it is not strictly required for biosynthesis of the APE_Ec_-carboxylic acid moiety. We did not detect any APE-related metabolites in the remaining 9 of the 18 mutant strains (Fig. S3). This was surprising for Δ*apeM* (transporter) since that strain’s extract retained a low level of yellow pigmentation (Fig. S2). In summary, except for *apeR*, all genes in the BGC seem required for full assembly of the APE_Ec_-carboxylic acid or proper attachment to its membrane anchor.

**Figure 4.**
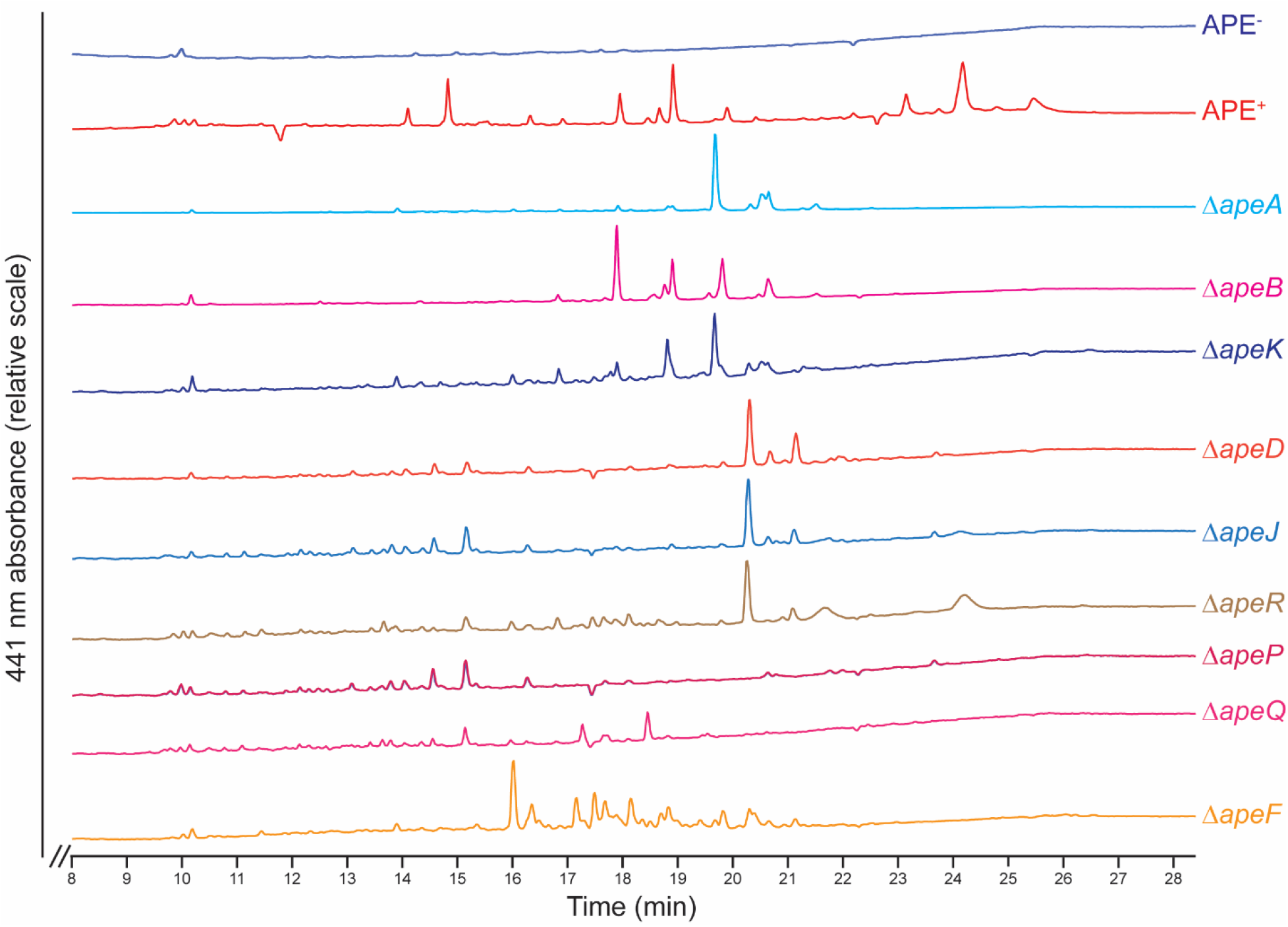
HPLC traces of *E. coli* Δ*ape* strain crude extracts. The APE^+^ extract (red) contained three clusters of APE-related peaks (around 14.5 min (57.5% acetonitrile (ACN)), 19 min (80% ACN) and 24 min (100% ACN)) which are not present in the APE^-^ strain (dark blue). The primary APE lipid eluting at 100% ACN only occurs in the APE^+^ and Δ*apeR* extracts. The Δ*apeA*, Δ*apeB* and Δ*apeK* strains produce a comparable set of immature APE carboxylic acids (early intermediates prior to initiation of the attachment process). Δ*apeD*, Δ*apeJ* and Δ*apeR* produce late pathway intermediates, which likely consist of APE carboxylic acids that are impaired in their proper attachment to the final outer membrane lipid. Deletion of the *apeF* ACP (orange) results in the formation of a series of yellow pigmented intermediates, indicating it is not strictly required for biosynthesis of the APE carboxylic acid moiety. HPLC traces for all Δ*ape* strains that did not have significant 441 nm absorbance are shown in Fig. S3. HPLC conditions used a gradient of ACN in 0.1% trifluoroacetic acid (TFA) water, initiating with a hold at 100% water for 3 min, followed by gradual increase from 0% to 100% ACN between 3 min and 23 min, and ending in a 100% ACN hold. Detection was at λ = 441nm.

### APE_Ec_ confers increased protection from oxidative, but not nitrosative challenge

Inspired by studies showing that staphyloxanthin deletion mutants of *S. aureus* are more susceptible to ROS [18, 29], we hypothesized that APEs fulfill a similar role for their Gram-negative producers. To test for protection against acute ROS stress, we compared APE^+^ and APE^-^ *E. coli* resilience in a hydrogen peroxide (H_2_O_2_) survival assay (Fig. 5A). Bacteria from an overnight liquid culture were challenged with incremental concentrations of H_2_O_2_ for one hour, after which the reaction was quenched by addition of excess catalase. Dilutions of the reaction mixtures were plated to count colony forming units and percent survival was calculated relative to the phosphate buffered saline (PBS) control incubated without added H_2_O_2_. At higher H_2_O_2_ concentrations, the APE^+^ strain had a significant fitness advantage compared to APE^-^. Our data suggest that APE_Ec_ expression can protect bacteria against acute ROS stress, such as from an oxidative burst.

**Figure 5.**
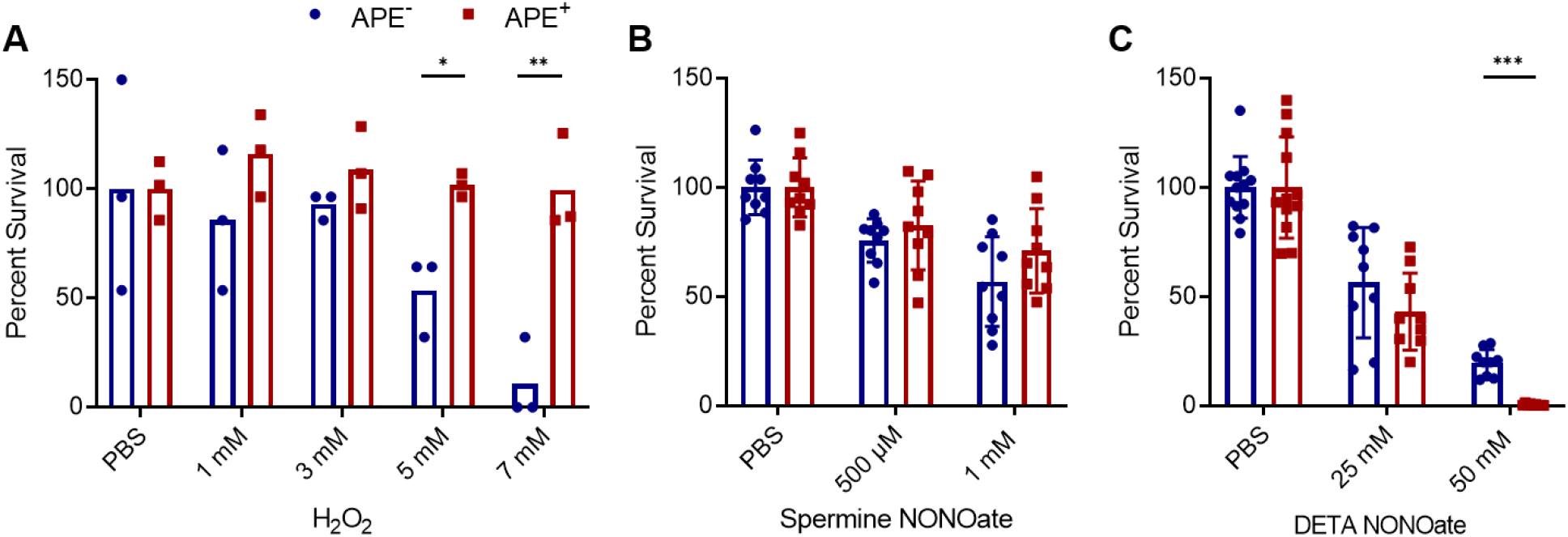
APE_Ec_ expression protects from oxidative, but not from nitrosative challenge. **A)** Percent survival upon exposure to different concentrations of hydrogen peroxide (H_2_O_2_) is calculated from colony forming units after a 60-minute H_2_O_2_ challenge for the APE_Ec_ expressing (APE^+^, red) versus control strain (APE^-^, blue) relative to the PBS control. The reactions were stopped by adding an excess of catalase. **B)** APE_Ec_ expression does not mediate protection from reactive nitrogen challenge. Percent survival is calculated from colony forming units after a 1 – 2 hour spermine NONOate challenge for the APE_Ec_ expressing (APE^+^) versus control strain (APE^-^) relative to the PBS control. **C)** Percent survival of APE^+^ and APE^-^ strains after 1-2 hours of DETA NONOate challenge. PBS = phosphate buffered saline (n=3-5 repeats; error bars represent SD; \**p*<0.05, \*\**p*<0.01 and \*\*\**p*<0.001 by unpaired t-test with Welch’s correction).

Given that in the context of infection ROS and RNS are both important mediators of immune response, we tested if APE producing bacteria were also protected from nitric oxide (NO) exposure. Bacteria from mid-log liquid cultures were dosed with NO donating NONOate compounds under aerobic conditions (Fig. 5B, C). Dilutions of the reaction mixtures were plated to count colony forming units and percent survival was calculated relative to the phosphate buffered saline (PBS) control incubated without NONOate. We did not observe a significant survival difference upon challenge with spermine NONOate (Fig. 5B). However, the APE^+^ strain had a notable decrease in survival relative to the APE^-^ strain when challenged with a supraphysiological level of 50 mM DETA NONOate (Fig. 5C). Granted, there was no discernable difference between APE^+^ and APE^-^ strains at the lower concentration. Taken together, these data suggest that contrary to the ROS assay, expression of APE_Ec_ does not confer a survival advantage to NO exposure.

### Expression of the APE_Ec_ BGC induces biofilm formation

We observed that APE^+^ bacteria form aggregates when grown in liquid culture and we hypothesized that the presence of APE_Ec_ in the outer membrane affects biofilm formation. To test this, we assessed the biofilm formation capacity of *E. coli* APE^-^ and APE^+^ in an *in vitro* biofilm assay. The individual strains were grown in non-treated, flat-bottom plates for 48 hours and subsequently stained with crystal violet and quantified by measuring optical absorbance levels at 595 nm. We observed a marked increase in biofilm formation for the APE^+^ strain (Fig. 6). Next, we set out to determine the contribution to biofilm formation for the individual genes within the APE_Ec_ BGC. Only the Δ*apeE* (ACP) and Δ*apeN* (LolB-like protein) strains showed a significant increase in biofilm formation when compared to APE^-^ (Fig. S4). Since APE_Ec_ biosynthesis is completely abolished in Δ*apeE* (Fig. S3), these data suggest that biofilm formation is not directly caused by APE biosynthesis. Instead, APE expression might cause alterations in regulatory cascades or cell envelope composition that result in increased biofilm formation.

**Figure 6.**
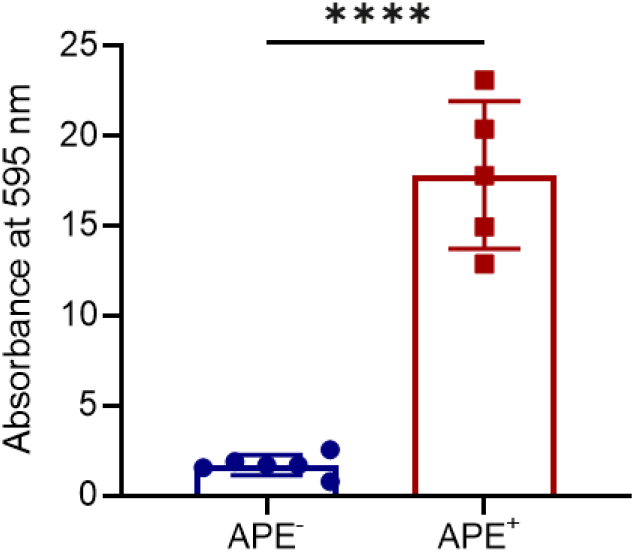
APE_Ec_ expression increases *E. coli* biofilm formation. Biofilms were grown in non-treated multi-well plates for 48 hours and subsequently stained with crystal violet. Absorbance at 595 nm was measured and normalized to APE^-^ values (n=3 repeats; error bars represent SD; \*\*\*\**p*<0.0001 by unpaired, two-tailed t-test).

To further characterize the biofilm structure and formation process over time, we visualized the biofilms at 48 and 72 hours using confocal fluorescence microscopy. This confirmed that the APE^+^ strain forms a robust and homogenous biofilm after 48 hours, in comparison to the disperse attachment observed for APE^-^ (Fig. 7A). However, after 72 hours, there is no discernable difference between APE^+^ and APE^-^ strains by confocal microscopy. In order to further investigate this finding, we acquired z-stacked images to assess the homogeneity and density of the respective biofilms. Three-dimensional rendering of these images shows that APE_Ec_ expression results in a much thicker biofilm of ~60 μm, compared to the ~29 μm thick APE^-^ aggregates (Fig. 7B). At 72 hours, the APE^-^ biofilm becomes more homogenous in structure, but at ~46 μm remains thinner than the denser ~53 μm APE^+^ biofilm. These observations were further quantified over time and showed that at 24 and 48 hours, APE^+^ biofilms are on average 2.5 times more voluminous than the APE^-^ counterparts (Fig. 7C). In conclusion, APE_Ec_ expression increases the rate of *E. coli* adherence and biofilm formation, resulting in a thicker and denser structure.

**Figure 7.**
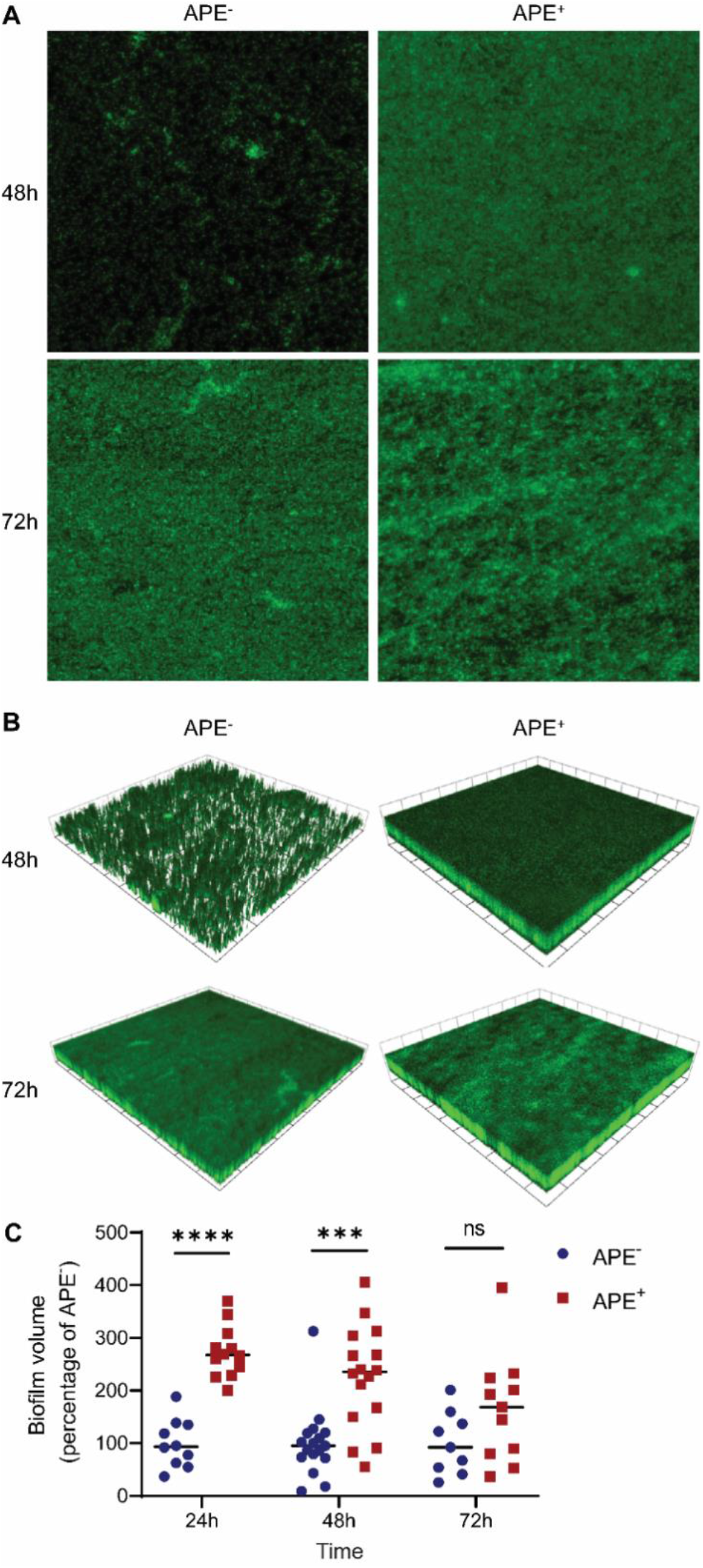
APE_Ec_ expressing *E. coli* more rapidly form thicker and denser biofilms. **A)** Confocal fluorescence microscopy images of APE^-^ and APE^+^ *E. coli* biofilms that were grown in non-tissue culture treated multi-well plates for 48 hours and 72 hours. Samples were fixed, stained with SYTO 9, and imaged using the 63× objective. **B)** Three-dimensional rendering of the z-stacked images for the APE^-^ and APE^+^ biofilms shows that APE_Ec_ expression results in formation of a thicker and denser biofilm. 1 scale unit = 57.97 μm. **C)** The volume of biofilm produced by APE^+^ (red) is on average 2.5 times higher than for APE^-^ (blue) after 24 and 48 hours of growth. Volocity (Quorum Technologies) was used to measure biofilm thickness, render 3D-images and calculate biofilm volumes. (n=3 repeats; \*\*\**p*<0.001, \*\*\*\**p*<0.0001 by unpaired, two-tailed t-test).

## DISCUSSION

We report the first genetic characterization and functional analysis of an APE BGC from a medically relevant bacterial uropathogen. The core biosynthesis for APE_Xd_, the product of a related BGC from the nematode symbiont *X. doucetiae*, was elucidated in a recent herculean *in vitro* effort by Grammbitter *et al*. [11]. Nine enzymes from the APE_Xd_ BGC (ApeC, E, F, H, I, O, P, Q and R) were produced heterologously in *E. coli* and several of these were found to form stable heterocomplexes. The biosynthetic roles for these enzymes were characterized and the formation of APE_Xd_-ACP was reconstituted *in vitro*. Our genetic analysis of the *E. coli* CFT073 APE_Ec_ BGC complements this biochemical analysis and contributes important *in vivo* insights into the APE biosynthetic process, including the identification of functionally redundant core biosynthetic genes, as well as non-essential genes for cell envelope attachment and transport later in the pathway.

Among the mutants in the core biosynthetic genes, Δ*apeE* is APE^-^, while Δ*apeF* retains partial pigmentation (Fig. 2), indicating that this latter ACP might be involved in malonyl chain extension and that the deletion could be partially complemented by the ACP involved in *E. coli* fatty acid biosynthesis, AcpP [30]. This also suggests that ApeF does not load 4-HBA *in vivo*, making ApeE the preferred ACP used for loading the 4-HBA starter (Figs. 3, 4). This observation is in line with the prior *in vitro* reconstitution of the respective APE_Xd_ pathway, which showed that both ACPs could be loaded with 4-HBA, albeit at a much lower efficiency for ApeF [11]. Our deletion analysis indicates that this does not occur at sufficiently high levels for *in vivo* pathway activity. Moreover, the functional redundancy of ApeF is further illustrated by the observation that not all APE BGCs have a second ACP as is the case for xanthomonadin APE BGCs from *Xanthomonas spp*. [1, 12]. Deletion of *apeK*, the gene encoding a putative thioesterase, resulted only in partial loss of pigmentation. No *in vitro* hydrolysis of 4-HBA-CoA or ACP-APE intermediates was observed for this enzyme, and a proofreading activity for ACP loading was proposed [11]. Since APE_Ec_-carboxylic acid production wasn’t abolished *in vivo*, this suggests that the putative proofreading activity is not absolutely required for biosynthesis but could have an impact on yield.

Gram-negative lipid biosynthesis requires adenosine triphosphate (ATP) and therefore typically takes place at the inner leaflet of the cell membrane. Lipids that are destined for the outer membrane are subsequently transported across the periplasm by dedicated transport systems [27]. Several of the genes predicted to be involved in APE_Ec_ cell envelope attachment (*apeD, apeJ* and *apeK*) still produce reduced pigmentation due to accumulated intermediates upon knockout, suggesting the biosynthesis of the core APE_Ec_ moiety is not disturbed (Figs. 2, 4). We propose that APE_Ec_ is transferred to an anchor molecule by ApeD (lysophospholipid acyltransferase) and ApeJ (a glycosyltransferase / acyltransferase). This anchoring might be facilitated by ApeK thioesterase-mediated release of APE_Ec_ from its ACP. The APE_Ec_-lipid might be transported across the cytoplasmic membrane by ApeM (an RND family transporter, Fig. 8). ApeG is a membrane protein with a predicted COG4648 pyrophosphatase domain that could be involved in providing the energy to drive transport through ATP hydrolysis, in a similar fashion to the role of SecA during protein translocation. Gram-negative bacteria shuttle lipids across the periplasmic space when their hydrophobic acyl chains are shielded from the aqueous milieu. This shielding can happen either via vesicular trafficking or via a protein-assisted process [27]. Since ApeL and ApeN share weak homology to components of the ‘localization of lipoprotein’ (Lol) pathway and ApeA encodes for a lipoprotein, these proteins could be involved in localization and insertion of the APE_Ec_-lipid to the outer membrane [27, 31]. ApeGLMN homologs are conserved across different APE BGCs [1], suggesting they constitute dedicated transport machinery with low similarity to currently characterized translocation systems. At present, the identity of the molecule that covalently anchors APE_Ec_ in the outer membrane is still not known. Conserved domains in the two acyltransferases ApeD and ApeJ suggests that this molecular anchor could be a glycolipid, potentially lipid A or a derivative thereof. Alternatively, the presence of a predicted lipoprotein and Lol-like transport genes could imply that APEs might get localized to the outer membrane while serving as a lipid anchor for ApeA. During the revision of this manuscript, Grammbitter *et al*. solved the structure of the APE-containing lipid from *X. doucetiae* [32]. It is feasible that the APE_EC_ is attached to a similar type of lipid in the *E. coli* cell envelope.

**Figure 8.**
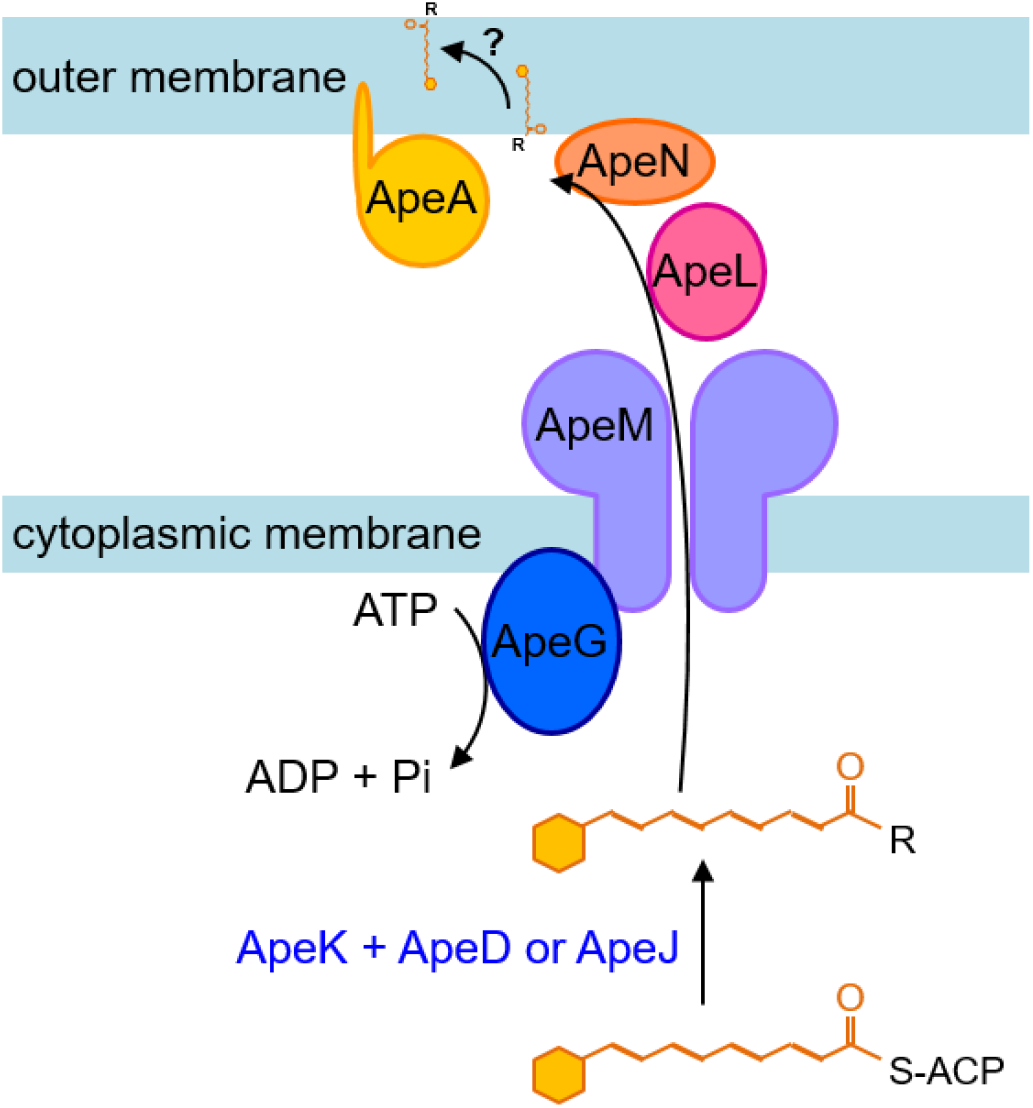
Schematic diagram for APE export and outer membrane localization. Following biosynthesis of the core APE_Ec_-carboxylic acid (Fig. 4), we propose that this entity is attached to a larger lipid anchor molecule (R) by ApeD (lysophospholipid acyltransferase) and ApeJ (a glycosyltransferase / acyltransferase). This attachment process might be assisted by the ApeK thioesterase-mediated release of APE_Ec_ from its ACP. The APE_Ec_-lipid might be transported across the cytoplasmic membrane by ApeM (an RND family transporter). The energy for this translocation could be provided by ATP hydrolysis catalyzed by ApeG (a predicted membrane-associated protein with a COG4648 pyrophosphatase domain). Since ApeL and ApeN share weak homology to components of the ‘localization of lipoprotein’ (Lol) pathway and ApeA encodes for a lipoprotein, these proteins could be involved in localization of the APE_Ec_-lipid to the outer membrane. Finally, the APE_Ec_ containing anchor lipid is potentially flipped to the outer leaflet of the outer membrane.

The generation of ROS constitutes a critical component of the initial line of defense by phagocytes against invading bacterial pathogens [20]. In turn, bacteria have developed several mechanisms to protect themselves from ROS damage. Enzymatic detoxification is an important mechanism employed to deal with an intracellular excess of ROS [33]. Three types of enzymes are key in this process: superoxide dismutases catalyze the dismutation of superoxide into O2 and H_2_O_2_, and catalases and peroxiredoxins detoxify H_2_O_2_ by enzymatic degradation [21]. As a complementary strategy, cells express polyenes in their envelope to form a protective shield that quenches ROS, thereby preventing their entry into the cell [18, 29]. One of the most well-known groups of pigments with a polyene chromophore that display antioxidant activity are the carotenoids. While their functional analogy to APEs in bacterial protection from ROS has been recognized [1, 4], carotenoids are produced by a biosynthetically distinct process. These lipophilic isoprenoids are assembled from isopentenyl pyrophosphate and dimethylallyl pyrophosphate building blocks by the subsequent action of a prenyl transferase, phytoene synthase and phytoene desaturase [34]. Examples of bacterial carotenoids that protect their producers from external ROS are the ‘golden’ pigment staphyloxanthin, produced by *S. aureus* (Fig. 1) and the red pigment deinoxanthin from *Deinococcus radiodurans* [18, 29, 35]. Amide-containing polyenes are a different family of pigments that are present in Gram-positive cell envelopes. These compounds are produced by a fatty-acid-like type II polyketide pathway more closely related to APE biosynthesis. The red amide-containing pigment granadaene is an ornithine rhamno-polyene produced by group B *Streptococcus* and it has been implicated in oxidative stress protection (Fig. 1, [17, 19]). A related amide-containing polyene was isolated from *Propionibacterium jensenii* [36] and a related BGC from commensal *Lactobacillus reuteri* was shown to produce a polyenic small molecule that stimulates the host aryl hydrocarbon receptor [37]. A potential role in bacterial protection from ROS remains to be investigated for the latter two examples. We challenged APE_Ec_ expressing bacteria with various concentrations of H_2_O_2_ and found that they significantly increased survival compared to the control strain (Fig. 5). Our results are corroborated by other studies showing that APEs protect from oxidative stress in a variety of different bacterial genera, including *Variovorax* [4], *Lysobacter* [6] and *Xanthomonas* [38–40]. Carotenoids and amide-containing polyenes present an interesting example of convergent evolution in Gram-positive bacteria and our results suggest an analogous function for APEs in Gram-negative bacteria.

Contrary to their ROS protective function, we did not observe a discernable survival advantage of APE_Ec_ expression upon RNS challenge. The differential susceptibility to 50 mM DETA-NONOate suggests more investigation is needed to understand if APE expression compromises respiratory metabolism flexibility and how this might impact *E. coli’s* competitive advantage in intestinal colonization [41]. Alternatively, our observation might be an artifact of the aerobic environment under which this experiment was completed. In addition, *E. coli* expresses a number RNS mediators such as flavohemoprotein (Hmp) which primarily functions in aerobic conditions, and flavorubredoxin (NorVW) and cytochrome c nitrate reductase (NrfA) which are only active under anaerobic or microoxic conditions [42]. These additional defenses may overshadow the contribution of aryl polyenes. In the context of infection where the phagosome enclosed bacteria will encounter both H_2_O_2_ and NO after the macrophage has activated iNOS, there is a prioritization of H_2_O_2_ mitigation by *E. coli* [43]. Whether aryl polyenes might provide some beneficial effect during acute exposure to ROS and RNS together may be better studied using an *in vitro* model.

Laboratory strains of *E. coli* K-12 are typically poor biofilm formers [44]. It was therefore surprising that heterologous expression of the APE_Ec_ BGC from the UPEC strain CFT073 in Top10 resulted in a robust increase in biofilm formation. There are various cell-envelope associated factors that contribute to *E. coli* adhesion to solid surfaces and biofilm formation, including type 1 and curli fimbriae, conjugative pili, autotransporters, cell surface polysaccharides, and capsules [44–46]. The mechanism by which APE_Ec_ expression contributes to the biofilm formation process remains to be determined. It is possible that APE expression contributes indirectly by activating regulatory cascades involved in biofilm formation. However, since APEs are likely part of a larger glycolipid molecule, it is feasible that the cell surface localization of this molecular entity structurally supports the polysaccharide matrix. Alternatively, APE expression could cause a change of the outer membrane that results in increased exposure of previously characterized cell-surface adhesion factors.

Uropathogenic *E. coli*, such as the APE_Ec_ parent strain CFT073, display a biofilm-like phenotype during urinary tract infection (UTI) [47]. After colonization of the urethra, bacteria ascend to the bladder where they can invade epithelial cells. The intracellular bacteria then replicate rapidly and form biofilm-like intracellular bacterial communities that can remain stable for multiple weeks, even after a standard course of oral antibiotic treatment has been completed. These bacteria can flux out of the epithelial cells in large numbers and cause the infection to resurge, and potentially ascend to the kidneys, thereby causing a more severe disease state such as pyelonephritis. The partly intracellular lifestyle of UPEC makes treatment challenging, since antibiotics cannot penetrate into the epithelial cells and the treatment regimen might be shorter than the time the bacteria persist intracellularly [48]. In addition, biofilm formation on indwelling devices, such as stents, implants, and catheters can form reservoirs of pathogenic bacteria thus leading to chronic infections. Catheter-associated urinary tract infections are the most common hospital acquired infections in the USA, and UPEC strains are the predominant cause [49]. Interestingly, overexpression of the transcriptional regulator TosR in *E. coli* CFT073 caused increased biofilm formation, as well as induced expression of the APE_Ec_ BGC [50], though no functional link between these two events has yet been established. Our future studies will focus on the antioxidant and biofilm-inducing roles of APEs in the context of *in vivo* bacterial pathogenesis.

Several examples exist within the Enterobacteriaceae, where APE BGCs are located on pathogenicity islands [1, 51, 52], suggesting they could be transmitted to other bacteria through horizontal gene transfer. This potential mobility is a major concern, especially given their roles in protection from oxidative stress and biofilm formation. It is therefore important to gain a fundamental understanding of APE biosynthesis and biology, in order to develop APE-targeting strategies for combating multidrug-resistant pathogens and preventing their biofilm formation.

Collectively, our data demonstrate the importance of studying the *in vivo* production of APEs, highlighting key differences not appreciable using elegant *in vitro* systems. Moreover, we have demonstrated a potential role for APEs in the pathogenic function of Gram-negative Enterobacteriaceae, mediating resistance to oxidative stress and facilitating robust biofilm formation. Our work provides essential molecular detail that could advance efforts to develop much-need therapeutic strategies to treat pathogenic *E. coli*.

## METHODS

### General methods, bacterial cultures and growth media

*E. coli* cultures were grown as liquid or solid agar media, using a Luria-Bertani (LB) or Mueller-Hinton base. Incubation was at 37°C, except for the strains carrying a temperature sensitive ori plasmid that required 28°C incubation. Antibiotic concentrations used for selection and maintenance were carbenicillin (100 μg/ml), kanamycin (50 μg/ml) and apramycin (50 μg/ml). Plasmid DNA isolation was performed with a standard kit according to the manufacturer’s instructions (Qiagen). General microbiological manipulations, PCR amplification, cloning and transformation procedures were as described previously [53]. Enzymes were purchased from New England Biolabs, and chemicals from MilliporeSigma or Fisher Scientific, unless otherwise stated. Oligonucleotide primers were obtained from Integrated DNA Technologies. Statistical analyses were performed as indicated in each figure legend, using GraphPad Prism version 9.0.0 (GraphPad Software).

### In-frame scarred mutant construction

The bacterial strains, primers and plasmids used or generated in this study are listed in Tables S1-3. Individual genes in the APE_Ec_ BGC were replaced with an 81 bp in-frame scar sequence in a two-step PCR-generated recombineering process [28]. The procedure is illustrated schematically in Fig. S1 for *apeK*, other genes were replaced in an analogous manner. pJC121 is the parent plasmid that was used for generating all Δscar derivatives ([1], Table S3). Briefly, in the first step, a targeting cassette is generated by PCR amplification of the FRT sites that flank the apramycin resistance gene from pIJ773. The primers used for amplification contain large (on average 58 nt) 5’ tails that are homologous to the up and downstream regions of the gene to be deleted (Table S2). The pIJ121 parent vector and targeting cassette are introduced into *E. coli* BW25113 with pIJ790 (a λRED recombination plasmid) and the genes of interest are replaced with the apra^R^ marker via double homologous recombination, yielding the respective Δapra^R^ constructs for each gene (even numbered plasmids pJC130-164 in Table S3). In the second step, the resistance markers are removed by introducing the Δapra^R^ plasmids into *E. coli* BT340. Expression of Flp recombinase by this strain causes recombination of the FRT sites that flank the apra^R^ marker, leaving an 81 bp scar sequence. The resulting in-frame scarred mutant plasmids can be identified based on their apramycin sensitivity (odd numbered plasmids PJC131-165 in Table S3).

### Analytical profiling of APE-related metabolites

A single colony of the APE^-^, APE^+^ and different Δape *E. coli* strains was used to inoculate 50 ml liquid cultures in LB + kanamycin (50 μg/ml). Liquid cultures were incubated for 24 hours at 37°C (250 rpm, in the dark) prior to harvesting the cell pellets by centrifugation (3500 g, 10 min). The growth medium was decanted and the cell pellets were washed twice by resuspension in dH_2_O followed by centrifugation. Cell pellets were extracted once with 25 ml acetone and then twice with 25 ml acetone:methanol (2:1 vol/vol). The extracts were filtered over paper and combined. Next, organic solvent was removed by a rotary evaporator. The dried samples were reconstituted in MeOH and passaged over a silica column prior to HPLC analysis. HPLC conditions used a gradient of acetonitrile in 0.1% trifluoroacetic acid water: Initiating with a hold at 100% water for 3 min, followed by gradual increase from 0% to 100% ACN between 3 min and 23 min, and ending in a 100% ACN hold for 3 minutes, followed by a 1 min re-equilibration to 100% aqueous phase and 3 min hold at 100% aqueous. Detection was at λ = 441nm.

### H_2_O_2_ survival assay

For H_2_O_2_ survival assays, a starter culture was grown overnight in LB, cells were pelleted and resuspended in PBS at OD_600_ = 1. Equal volumes of H_2_O_2_ (in PBS) were added with final concentrations ranging from 0 to 7 mM. Reactions were incubated at room temperature for 1 hour, quenched by addition of excess catalase and serially diluted prior to plating onto LB agar. Colony forming units were counted after overnight incubation at 37°C and percent survival was calculated relative to the PBS only control.

### NONOate survival assays

For the NONOate survival assays, a starter culture was grown for 4hrs at 37°C, 220 rpm. The cultures with similar starting OD readings were spun down and washed with PBS. The pellet was resuspended in PBS to a final OD_600_ of 0.3. Equal volumes of DETA NONOate (Cayman, 82120) or spermine NONOate (Cayman, 82150) were diluted in PBS and added to the bacteria. The final concentrations of the reactions were 25 and 50 mM for DETA NONOate, and 0.5 or 1 mM for spermine NONOate with PBS controls. Reactions were incubated for 1 hour (DETA NONOate) or 2 hours (spermine NONOate) and serially diluted prior to plating onto LB agar. Colony forming units were counted after overnight incubation at 37°C and percent survival was calculated relative to the PBS only control.

### *E. coli* biofilm assays

An overnight culture of APE^+^ and APE^-^ *E. coli* was adjusted to OD_600_ = 1, and 10 μl was used to inoculate a total of 1 ml fresh medium in flat-bottom, non-tissue culture treated plates. Biofilms were grown by static incubation at 37°C for 24, 48 or 72 hours. All steps were performed with care to not disturb the biofilms. After removing the supernatant, cells were fixed for 30 minutes with 250 μl Bouin’s solution (Sigma). The fixing solution was removed and cells were washed twice with 250 μl sterile H_2_O prior to staining for quantification or confocal imaging. For quantitative staining, fixed samples were submerged in 250 μl of a 0.1% v/v crystal violet solution for 15 minutes. The staining solution was removed and samples were washed twice with 250 μl sterile H_2_O. The remaining material was dissolved in 300 μl of 95% v/v ethanol, and quantified by measuring absorbance at 595 nm. Fixed biofilm samples that were processed for confocal imaging were stained with 250 μl of SYTO 9 green fluorescent dye. Confocal laser scanning microscopy image acquisition was performed using a Leica TCS SP5, using the 20× objective. Volocity (Quorum Technologies) was used to measure biofilm thickness and to produce three-dimensional rendered images by combining z-stacked images that were recorded at a z-step of 0.25 μm. Biofilm volumes were calculated by using the Volocity measurements and the software’s “find object” features.

## ACKNOWLEDGEMENTS

We thank Dr. John W Peterson and Dr. Judith A Drazba of the LRI imaging core for assistance with the biofilm imaging.

## Author contributions

Conceptualization and design: IJ, JC. Investigation: IJ, LJO, RLM, EAM, AK, KBS, NN, JC. Figures: IJ, LJO, RLM, JC. Writing of the original draft: IJ, JC. Writing, review and interpretation: IJ, LJO, RLM, PPA, JMB, JC.

## Materials and correspondence

Please address all correspondence and material requests to JC at

