## Supplemental information for "Identification of essential genes for *Escherichia coli* aryl polyene biosynthesis and function in biofilm formation"

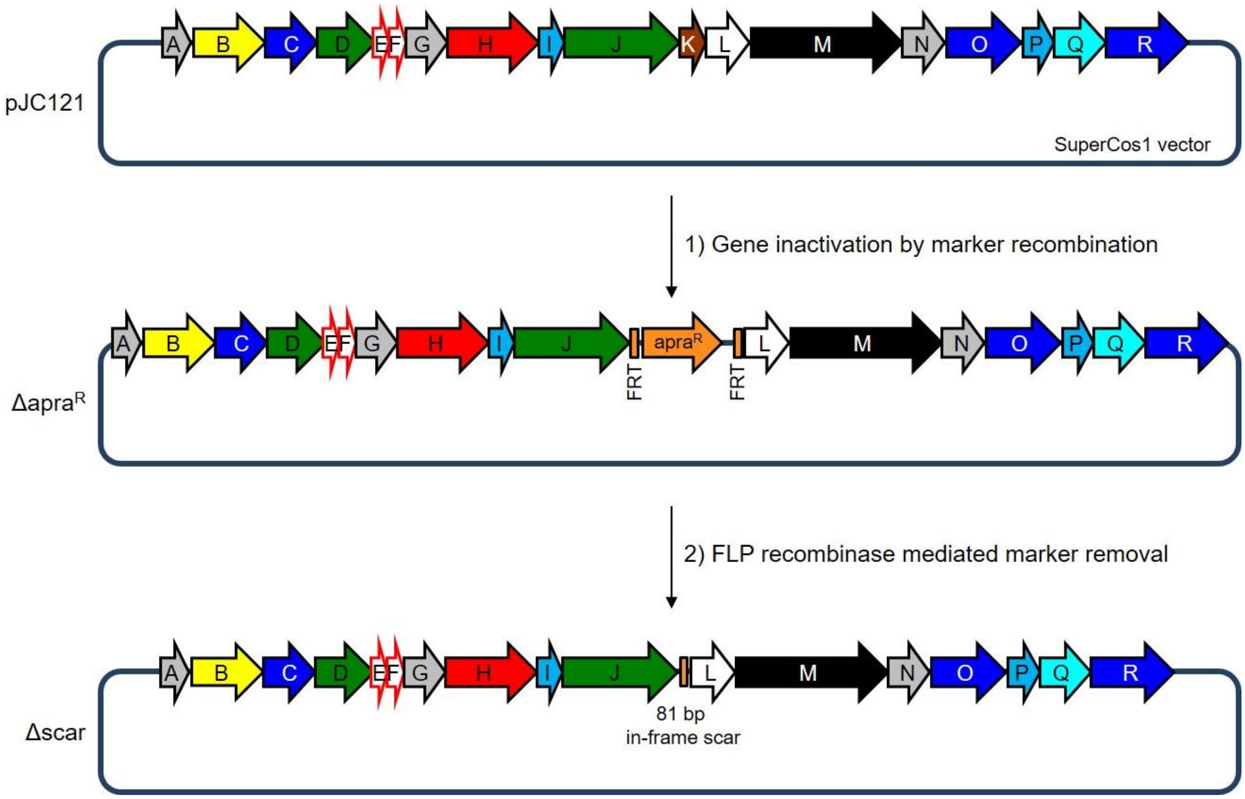


**Supplementary Figure 1. Schematic overview of scar mutagenesis procedure.** Individual genes in the $APE_{Ec}$ BGC were replaced with an 81 bp in-frame scar sequence in a two-step PCR-generated recombineering process [28]. This is exemplified by *apeK* deletion, other genes were replaced in an analogous manner. **Step 1)** The pIJ121 parent vector and a gene-specific targeting cassette are introduced into an *E. coli* strain expressing the λRED recombination machinery, and the gene of interest is replaced with the apra$^R$ marker via double homologous recombination, yielding Δapra$^R$ plasmids for each gene (even numbered plasmids pJC130-164 in Table S3). **Step 2)** The resistance marker is removed via Flp mediated recombination of the apra$^R$ flanking FRT sites, yielding in-frame Δscar plasmids (odd numbered plasmids PJC131-165 in Table S3).

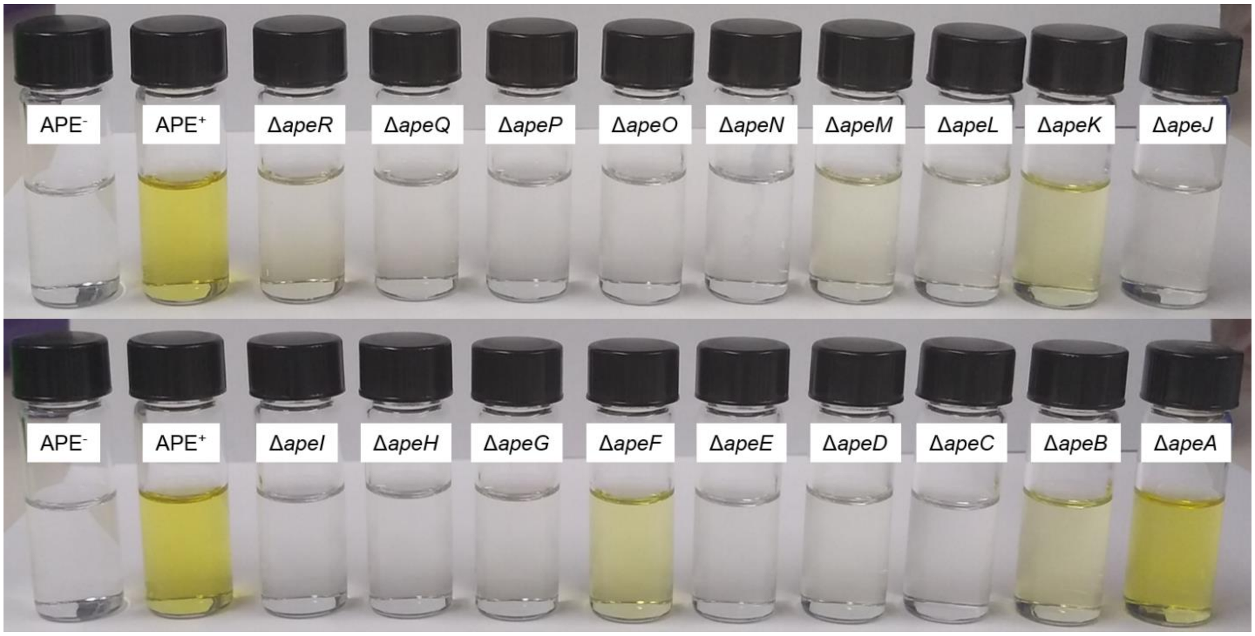


**Supplementary Figure 2. Crude cell extracts of the different *E. coli* Δ*ape* strains.** Cell pellets from the $APE^-$, $APE^+$ and various Δ*ape* mutants (Fig. 2) were extracted, filtered over paper, and evaporated to dryness prior to reconstitution in MeOH.

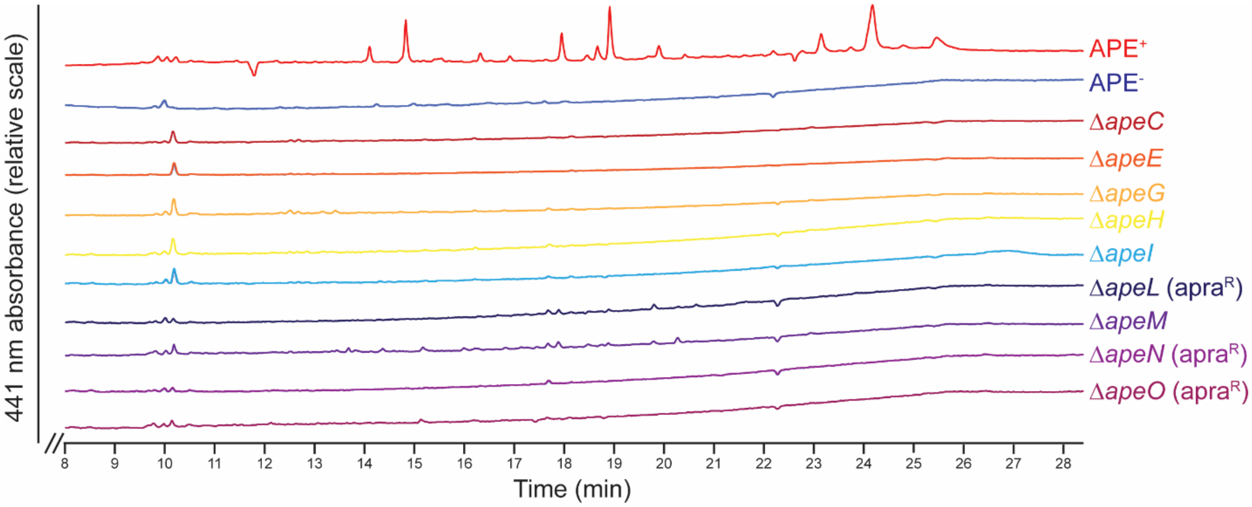


**Supplementary Figure 3. HPLC traces of *E. coli* Δ*ape* strain crude extracts that did not have significant 441 nm absorbance.** These strains comprise mostly the genes involved in biosynthesis of the core APE$_{Ec}$-carboxylic acid moiety (*apeC*, *apeE*, *apeHI* and *apeO*), as well as some genes predicted to be involved in transport (*apeG* and *apeLMN*). The APE$^+$ extract (red) and APE$^-$ strain (dark blue) are reproduced here from Fig. 3 for reference. Samples were run under the same HPLC conditions (a gradient of ACN in 0.1% trifluoroacetic acid (TFA) water, initiating with a hold at 100% water for 3 min, followed by gradual increase from 0% to 100% ACN between 3 min and 23 min, and ending in a 100% ACN hold). Detection was at $\lambda$ = 441nm.

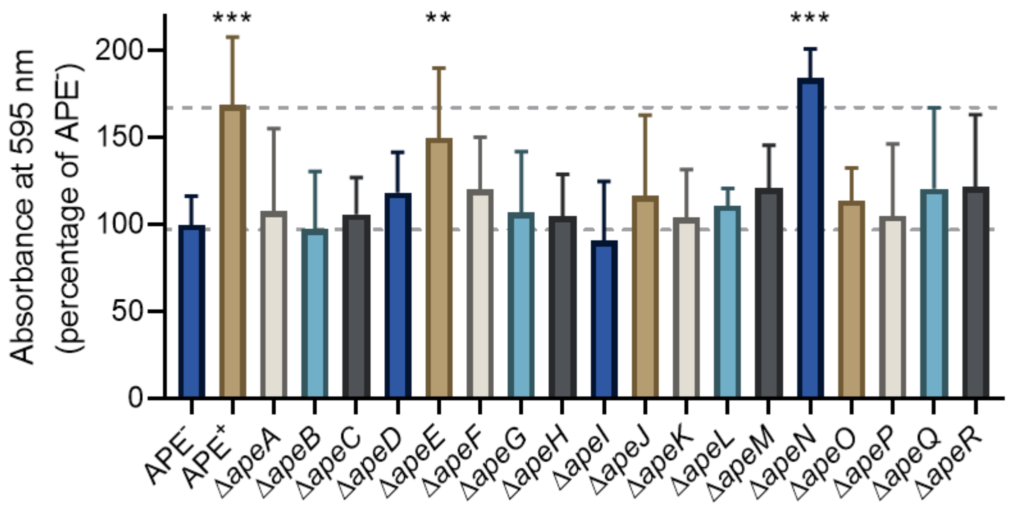


**Supplementary Figure 4. Crystal violet quantification of biofilms formed by the different Δ*ape* strains.** The APE$^+$, Δ*apeE* and Δ*apeN* strains showed a significant increase in biofilm formation compared to APE$^-$. Biofilms were grown in non-treated multi-well plates with MH medium for 48 hours and subsequently stained with crystal violet. Absorbance at 595 nm was measured and normalized to APE$^-$ values (n=3 repeats; error bars represent SD; **$p<0.01$, ***$p<0.001$ by unpaired, two-tailed t-test).

**Supplementary Table 1. Strains used in this study.**

| Strain | Description | Reference |
|---|---|---|
| CFT073 | uropathogenic E. coli strain, serotype O6:H1:K2 | [54] |
| Top10 | E. coli cloning strain | Invitrogen |
| BW25113 | E. coli K-12 derivative: ΔaraBAD, ΔrhaBAD | [55] |
| BT340 | E. coli DH5α/pCP20 | [56] |

**Supplementary Table 2. Primers used in this study.**

| Primer | Sequence (5'→3') |
|---|---|
| apeA F | AATTATATAATTGTATTGCATACATAAGAGGCACTAATGATTCCGGGGATCCGTCGACC |
| apeA R | ATACTATCGGGCCATTCCAGCCCGATATCAGCATCACTATGTAGGCTGGAGCTGCTTC |
| apeB F | GCCATCACCGAAGCACAGCGGATTGCCTTTGCTCCTATGATTCCGGGGATCCGTCGACC |
| apeB R | GCGCGCCCGCGGTAAACCAAAATTATTAAGTGACGATTATGTAGGCTGGAGCTGCTTC |
| apeC F | CACTTAATAATTTTGGTTTACCGCGGGCGCGCGTTGATGATTCCGGGGATCCGTCGACC |
| apeC R | TCATCTGCGGCTCCAGCGCCATTGCACGCGCTCGCCAGGTGTAGGCTGGAGCTGCTTC |
| apeD F | CGTGCAATGGCGCTGGAGCCGCAGATGAGTAAGCTGATGATTCCGGGGATCCGTCGACC |
| apeD T | TTAATTTCCAGATAAAGCGCTTGCATTATTTATTCCTGATGTAGGCTGGAGCTGCTTC |
| apeE F | TAAATAATGCAAGCGCTTTATCTGGAAATTAAAAACCTCATTCCGGGGATCCGTCGACC |
| apeE R | TGGTTTGTTGGTCTGTCATGATGGTTTTCCTCAGGCACGTGTAGGCTGGAGCTGCTTC |
| apeF F | TTTCATTGCTGCGCAACGTGCCTGAGGAAAACCATCATGATTCCGGGGATCCGTCGACC |
| apeF R | AGCGAGCGAATGCCCGACATCAAATTAGCCTTCTTGTAGTGTAGGCTGGAGCTGCTTC |
| apeG F | AGGCTGTAGAACGCCTGCTACAAGAAGGCTAATTTGATGATTCCGGGGATCCGTCGACC |
| apeG R | GTGAAAGAGAGAGAGTCTGATTCATGGTGTTTTCCGTTTTGTAGGCTGGAGCTGCTTC |
| apeH F | ACGTCGTAAGATGATCAAACGGAAAACACCATGAATCAGATTCCGGGGATCCGTCGACC |
| apeH R | CATGAAATAACTCCTGTAATTGCGCATAGACACGCTTATTGTAGGCTGGAGCTGCTTC |
| apeI F | AATTACAGGAGTTATTTCATGATACGCCATGAAATTGAGATTCCGGGGATCCGTCGACC |
| apeI R | GGTTGTAGCAGGGGATCAACACGCAGGGAGAAAAGTTTATGTAGGCTGGAGCTGCTTC |
| apeJ F | GCGATGATGCCGGGCGTGCTGGCGCGTCTTAAGCCATTTATTCCGGGGATCCGTCGACC |
| apeJ R | GGTAAATCGGGGATCGTTAAGCACCCTTTACTCCTTGTCTGTAGGCTGGAGCTGCTTC |
| apeK F | TGGCAACTGCCGGAAATTCAGGACAAGGAGTAAAGGGTGATTCCGGGGATCCGTCGACC |
| apeK R | GCGCCAGCAGCGGTAAAAATTTCATGGTTTGACTCCCATTGTAGGCTGGAGCTGCTTC |
| apeL F | TATTCTGTTTGAACGCATGGGAGTCAAACCATGAAATTTATTCCGGGGATCCGTCGACC |
| apeL R | GGCAAAACGTTGGTGTTCGTCATCGGTCAGTTGCGCTGGTGTAGGCTGGAGCTGCTTC |
| apeM F | CCGATGACGAACACCAACGTTTTGCCGCCCAGTAAACGCATTCCGGGGATCCGTCGACC |
| apeM R | CTCGCCAGAATGTGGATTTGATCATTTTTTTGTTCTCTTTGTAGGCTGGAGCTGCTTC |
| apeN F | GCTGGCGATGCCCGATAAAAAGAGAACAAAAAAATGATCATTCCGGGGATCCGTCGACC |
| apeN R | GCGGAAATATAAATCATATCAGTCACCTAAGTATTGAATTGTAGGCTGGAGCTGCTTC |
| apeO F | AATACCACATCACCATTCAATACTTAGGTGACTGATATGATTCCGGGGATCCGTCGACC |
| apeO R | CGCCGGGGGATAAATAGTGGCTCACGAAACCCTCCCGAGTGTAGGCTGGAGCTGCTTC |
| apeP F | TAACGCCAGCATTCTGCTCGGGAGGGTTTCGTGAGCCACATTCCGGGGATCCGTCGACC |
| apeP R | TAACCAGAACTGAACGACTCATCAGGACGCTCCTTGTTGTGTAGGCTGGAGCTGCTTC |
| apeQ F | ACTTACCACCCTGTTTCAACAAGGAGCGTCCTGATGAGTATTCCGGGGATCCGTCGACC |
| apeQ R | TAATCACTACGCGACGTGTCATAACATCCCTCCATTGATTGTAGGCTGGAGCTGCTTC |
| apeR F | CCGCCAGGTTATTTCCATCAATGGAGGGATGTTATGACAATTCCGGGGATCCGTCGACC |
| apeR R | GAATCACCACATGGGATGTCCATGTGGTTGTATAACTCATGTAGGCTGGAGCTGCTTC |

**Supplementary Table 3. Plasmids used in this study.**

| Plasmid | Description | Reference |
|---|---|---|
| SuperCos1 | cosmid vector (KmR ApR) | Agilent Inc. |
| pIJ773 | pBS SK+ with cassette P1-FRT-oriT-aac(3)IV-FRT-P2 | [28] |
| pIJ790 | λ-RED (gam, bet, exo), cat, araC, rep101ts | [28] |
| pJC121 | SuperCosI::CFT073-ape cluster (c1186-c1204) | [1] |
| pJC130 | pJC121 ΔapeA::apraR | present study |
| pJC131 | pJC121 ΔapeA::scar | present study |
| pJC132 | pJC121 ΔapeB::apraR | present study |
| pJC133 | pJC121 ΔapeB::scar | present study |
| pJC134 | pJC121 ΔapeC::apraR | present study |
| pJC135 | pJC121 ΔapeC::scar | present study |
| pJC136 | pJC121 ΔapeD::apraR | present study |
| pJC137 | pJC121 ΔapeD::scar | present study |
| pJC138 | pJC121 ΔapeE::apraR | present study |
| pJC139 | pJC121 ΔapeE::scar | present study |
| pJC140 | pJC121 ΔapeF::apraR | present study |
| pJC141 | pJC121 ΔapeF::scar | present study |
| pJC142 | pJC121 ΔapeG::apraR | present study |
| pJC143 | pJC121 ΔapeG::scar | present study |
| pJC144 | pJC121 ΔapeH::apraR | present study |
| pJC145 | pJC121 ΔapeH::scar | present study |
| pJC146 | pJC121 ΔapeI::apraR | present study |
| pJC147 | pJC121 ΔapeI::scar | present study |
| pJC148 | pJC121 ΔapeJ::apraR | present study |
| pJC149 | pJC121 ΔapeJ::scar | present study |
| pJC150 | pJC121 ΔapeK::apraR | present study |
| pJC151 | pJC121 ΔapeK::scar | present study |
| pJC152 | pJC121 ΔapeL::apraR | present study |
| pJC154 | pJC121 ΔapeM::apraR | present study |
| pJC155 | pJC121 ΔapeM::scar | present study |
| pJC156 | pJC121 ΔapeN::apraR | present study |
| pJC158 | pJC121 ΔapeO::apraR | present study |
| pJC160 | pJC121 ΔapeP::apraR | present study |
| pJC161 | pJC121 ΔapeP::scar | present study |
| pJC162 | pJC121 ΔapeQ::apraR | present study |
| pJC163 | pJC121 ΔapeQ::scar | present study |
| pJC164 | pJC121 ΔapeR::apraR | present study |
| pJC165 | pJC121 ΔapeR::scar | present study |
